# Landscape configuration and community structure jointly determine the persistence of mutualists under habitat loss

**DOI:** 10.1101/2025.05.28.655631

**Authors:** Subhendu Bhandary, Klementyna A. Gawecka, Fernando Pedraza, Jordi Bascompte

## Abstract

Habitat loss poses a major threat to biodiversity. Its effects on ecological communities depend on the complex interplay between the landscape configuration — the patterns of connections between habitat patches, community structure — the patterns of interactions between species, and habitat loss patterns. Despite their individual importance, their joint effect on species persistence remains poorly understood. We explore how these three factors influence the persistence of empirical mutualistic communities. By employing spatially explicit metacommunity models, we find that landscapes with a heterogeneous distribution of connections between habitat patches exhibit high persistence under spatially-uncorrelated habitat loss but are highly vulnerable to spatially-correlated loss, where adjacent habitat patches are destroyed sequentially. Homogeneous landscapes with regularly arranged patches have lower persistence than heterogeneous landscapes but are more resilient to correlated habitat loss. The metacommunity nested structure of species interactions enhances persistence, with varying magnitude depending on landscape configuration and the patterns of habitat loss. These results can guide conservation strategies by identifying landscape and community features that promote species persistence.

## Introduction

Habitat loss is one of the leading drivers of biodiversity decline, with far-reaching effects on ecological communities and their functioning (Chase et al., 2020; Tylianakis et al., 2008). The loss of species can propagate through ecological communities, resulting in further extinctions of species and interactions (Memmott et al., 2004). In mutualistic communities, this equates to the loss of key ecosystem services such as pollination and seed dispersal (Bascompte and Jordano, 2007). Importantly, habitat loss also reshapes spatial connectivity, fragmenting landscapes and reducing opportunities for recolonization (Kindlmann and Burel, 2008; Turner and Ruscher, 1988), which can push systems toward tipping points (Rockström et al., 2009; Scheffer et al., 2001). Understanding how species persistence is shaped by habitat loss, the landscape, and ecological community structure is crucial for anticipating collapse and guiding conservation responses.

The landscape configuration influences metacommunity dynamics, dispersal processes, and overall system stability (Arancibia, 2024; Hanski, 1999; Urban and Keitt, 2001). Graph-theoretic approaches provide a useful framework for characterizing landscapes as networks, where habitat patches are represented as nodes connected by dispersal pathways. The structure of these spatial networks determines how readily species move across the landscape and respond to disturbance. For example, homogeneous networks where patches are arranged in a regular grid (hereafter “grid”) promote local stability through uniform connectivity but may limit large-scale dispersal (Hanski and Ovaskainen, 2000). Networks with patches connected randomly (hereafter “random”) offer moderate connectivity and are relatively robust to stochastic perturbations (Erdos et al., 1960). While the heterogeneous scale-free networks, which contain few highly connected hub patches and many poorly connected peripheral patches, are robust to random node loss but remain highly vulnerable to removal of the hubs (Barabási and Albert, 1999). Thus, the spatial pattern of habitat destruction further shapes extinction trajectories: spatially correlated loss can sever entire regions and connectivity clusters, whereas spatially uncorrelated loss disrupts patches more diffusely (Keitt et al., 1997). While previous studies have explored spatial dynamics or habitat loss effects independently, the interaction between landscape configuration, habitat loss patterns, and ecological processes remains poorly understood.

Mutualistic communities, such as those composed of plants and their pollinators or seed-dispersers, are structured non-randomly in a way that promotes species coexistence and buffers against collapse (Bascompte et al., 2003; Bastolla et al., 2009; Lever et al., 2014). Here too, network theory provides a useful tool for analyzing the patterns of interactions between species within communities. For example, nestedness — a common pattern where specialist species interact with subsets of generalist partners — enhances community persistence by enabling redundant interaction pathways and indirect facilitation (Saavedra et al., 2011; Thébault and Fontaine, 2010). Generalist species often act as anchors for community resilience, whereas specialists tend to be more prone to extinction (Hagen et al., 2012). However, empirical and theoretical studies on the persistence of mutualistic communities rarely consider landscape-scale processes (Guimarães Jr, 2020). In fragmented landscapes, mutualistic community dynamics unfold across spatially structured habitats. Yet, the combined role of landscape configuration and mutualistic community structures in driving extinction remains largely unexplored.

Although important progress has been made in understanding the persistence of metacommunities in fragmented systems (Fortuna et al., 2013; Gilarranz and Bascompte, 2012; Liao et al., 2016), previous models often treat habitat loss, landscape configuration, and community structures in isolation. For instance, many studies focus on a single species or use synthetic interaction networks without taking into account realistic mutualistic community structures (Gilarranz and Bascompte, 2012; Guo et al., 2022; Liao et al., 2020; Zhang et al., 2021). Others adopt spatially homogeneous models that ignore how dispersal constraints modulate landscape-scale dynamics (Liao et al., 2022; Shen et al., 2019). A more integrative framework is needed to understand the interplay between spatial and community dynamics under different habitat loss regimes.

Here, we investigate how species persistence is shaped by the interaction between landscape configuration, mutualistic community structure, and habitat loss patterns. We achieve this by developing a spatially explicit metacommunity model. We embed empirical mutualistic communities with varying structure within three types of spatial networks: grid, random, and scale-free, and simulate dynamics under spatially-correlated and uncorrelated habitat loss. By linking ecological network theory with spatial dynamics, our work offers mechanistic insights into biodiversity collapse under habitat loss and informs conservation strategies for fragmented landscapes.

## Methods

We modeled landscapes as spatially explicit unipartite networks, where habitat patches were represented as nodes and dispersal routes as links. Each landscape contained 2,500 patches and 5,000 links, ensuring an average of four connections per patch. Connectivity varied along a gradient of heterogeneity: from grid networks, where each patch had exactly four connections, to random networks with links assigned randomly, and scale-free networks characterized by a power-law distribution of links while maintaining an average degree of four (Figure 1).

**Figure 1.**
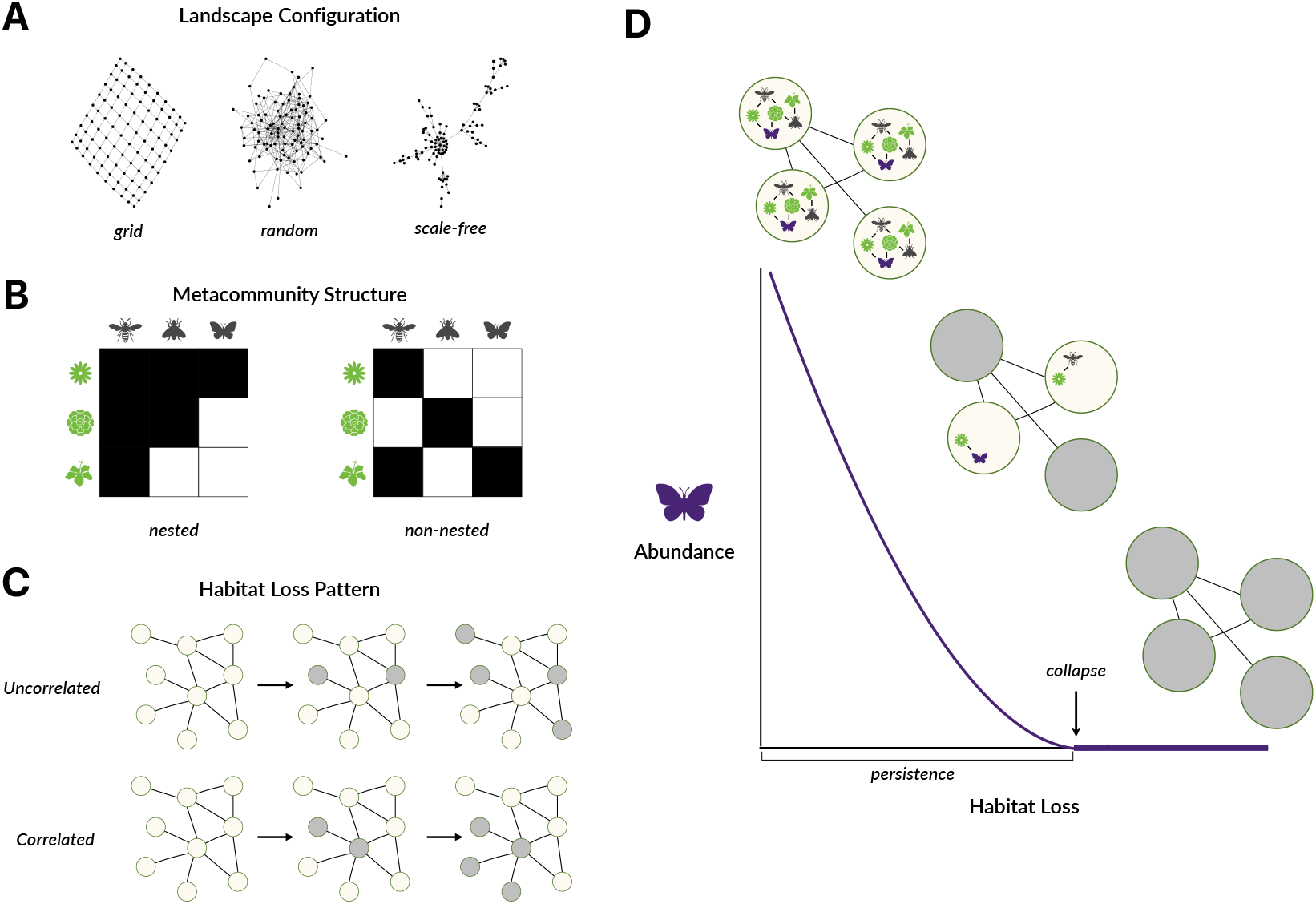
Schematic of our simulation treatments (A-C) and model output (D). We investigate the interplay between landscape configuration, mutualistic community structure, and habitat loss scenarios in shaping species persistence. (A) To study the effect of landscape structure, we consider networks of habitat patches with increasing heterogeneity in the distribution of connections per patch — from “grid” to “random” to “scale-free”. (B) We investigate the effect of mutualistic community structure by adopting empirical networks with varying nestedness — a pattern whereby a core group of generalist species interacts with each other, and extreme specialists interact with generalist species. (C) We simulate two scenarios of habitat loss. In the “uncorrelated” scenario, we destroy habitat patches in a random sequence, whereas in the “correlated” scenario, patches are lost in a clusters. (D) As habitat is destroyed, species transition from persistence state to extinction state.

We used 20 empirical mutualistic networks (10 plant-pollinator and 10 seed-dispersal) from the Web of Life database (www.web-of-life.es) (Fortuna et al., 2014) as metanetworks, which integrated all species and their interactions across the entire landscape (Fortuna et al., 2014). Within each patch, populations of plants and animals formed local networks which were subsets of the metanetwork. We selected empirical networks spanning a range of structural properties (Table T1 in Supplementary), particularly focusing on variation in network nestedness.

We measured network nestedness following the approach proposed by (Fortuna et al., 2019), which is based on the NODF (Nestedness metric based on Overlap and Decreasing Fill) metric (Almeida-Neto et al., 2008). This method quantifies nestedness by assessing the extent to which species with fewer interactions are linked to subsets of the partners of more connected species. It captures the average overlap between species interactions without penalizing networks where species have similar numbers of connections (as in NODF). To control for the effects of network size and connectance, we standardized the observed nestedness against expectations from a null model. Specifically, we generated 100 randomized networks per empirical network by preserving the marginal totals (row and column sums) and recalculated nestedness for each. We then computed the mean (*μ*) and standard deviation (*σ*) of nestedness from the randomized networks. Finally, we obtained the standardized nestedness z-score as:

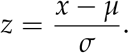

where *x* is the observed nestedness of the empirical network. This approach allowed us to compare nestedness across networks of different sizes and link densities.

At each time step, species faced a probability of extinction within the patches they occupied. Extinction probabilities were assumed to be uniform across species and patches. For the resource species (i.e., plants) and the mutualistic consumers (i.e., pollinators or seed dispersers), extinction probabilities were defined as (Gawecka and Bascompte, 2023):

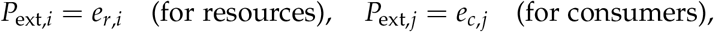

Where *e*_*r,i*_ and *e*_*c,j*_ are the intrinsic extinction rates of resource *i* and consumer *j*, respectively.

Colonization occurred between directly connected patches. For both resources and consumers, the probability of colonization increased with the number of neighboring patches and the presence of interaction partners (Gawecka and Bascompte, 2023):

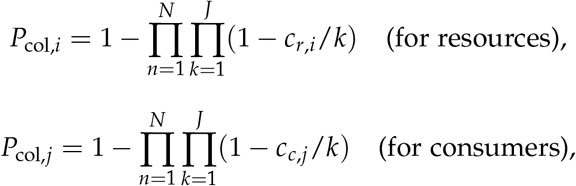

Where *c*_*r,i*_ and *c*_*c,j*_ are the intrinsic colonization rates of resource *i* and consumer *j*, respectively, *J* is the number of interaction partners of species *i*, and *N* is the number of neighboring patches that are suitable sources for colonization.

The system was simulated over 1,000 time steps to capture metacommunity dynamics, discarding the first 900 time steps to remove transient effects. Steady-state abundances were calculated as the mean number of patches occupied by each species over the final 100 time steps.

To investigate habitat loss effects, we incrementally removed habitat patches in 5% steps, starting from pristine landscapes (0% habitat loss) and continuing until all patches were destroyed. We modeled two habitat loss scenarios: (1) “correlated”, where destruction began at 25 (1%) randomly selected patches and extended to their connected neighbors, and (2) “uncorrelated”, where patches were destroyed randomly without regard for connectivity (Figure 1). After each increment, the system was simulated until reaching a new steady state.

We varied the extinction-to-colonization ratios for resources (*e*_*r,i*_/*c*_*r,i*_) and consumers (*e*_*c,j*_/*c*_*c,j*_) from 0 to 3 in steps of 0.3, keeping colonization rates fixed at *c*_*r,i*_ = *c*_*c,j*_ = 0.1 (Gilarranz and Bascompte, 2012) (Supplementary Fig. S1, S4, S5). These ratios capture a wide range of ecological dynamics, from populations with low extinction risk and high colonization potential to those with high extinction risk and limited colonization. This resulted in 441 unique parameter combinations, spanning a broad spectrum of demographic dynamics for both resources and consumers. Persistence probability was defined as the proportion of parameter combinations where the mean abundance of all species exceeded zero. This approach allows us to account for a wide range of extinction and colonization rates, thus generalizing our results. We quantified this persistence probability at each fraction of habitat loss.

All simulations were performed in Julia version 1.4.2 (Bezanson et al., 2017), and data visualization was performed in R version 3.6.2 (R Core Team, 2024).

## Results

The rate of decline of species persistence probability with habitat loss is highly dependent on the interplay between spatial landscape structure and habitat loss pattern (Fig.2). Grid landscapes, which have homogeneous connectivity, show lower persistence in pristine landscapes than the more heterogeneous random and scale-free networks. Yet, they exhibit the greatest robustness to habitat loss. In fact, under correlated habitat loss they enable the highest species persistence out of the three landscapes. In contrast, scale-free landscapes, characterized by highly connected hub patches, show the highest persistence probability at low habitat loss fractions. These networks are particularly resilient to uncorrelated habitat loss, maintaining persistence even under high levels destruction. However, under correlated habitat loss, their persistence declines very sharply, likely due to the targeted loss of key hub patches and the resulting cascade of extinctions. Random landscape configuration display intermediate behavior, with persistence probabilities declining steadily with habitat loss. They respond relatively similarly to both habitat loss patterns, suggesting lower sensitivity to how habitat is removed. In summary, landscapes with homogeneous connection between patches are resilient to spatially-correlated habitat loss, whereas heterogeneous ones exhibit high resilience to spatially-uncorrelated loss of patches.

We also find variability among mutualistic communities in their response to habitat loss in different landscapes (see the widths of boxplots in Fig. 2). High variability indicates that persistence depends on the structure of the community. At low habitat loss fractions, grid and random landscapes display greater variability in persistence than scale-free landscapes. As habitat loss progresses, grid and random networks show reduction in variability among communities, especially under uncorrelated habitat loss. In contract, scale-free landscapes exhibit an increase up to intermediate levels of habitat loss, followed by a reduction. Overall, these results highlight that the importance of community structure in driving persistence depends on the interaction between landscape configuration, habitat loss pattern, and the level of habitat destruction.

**Figure 2.**
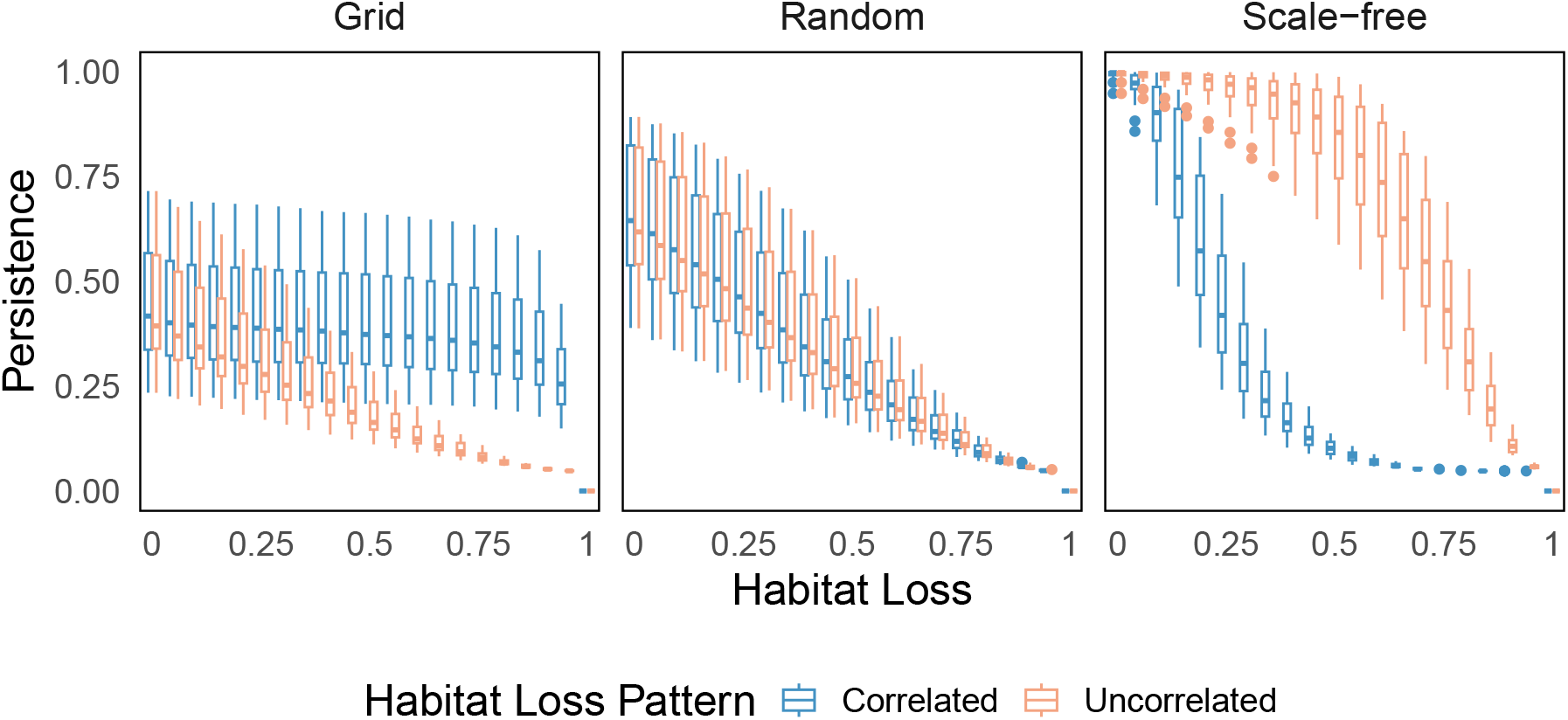
Species persistence probability under habitat loss. Persistence probability represents the proportion of extinction-to-colonization parameter combinations where species abundance remains above zero. Boxplots show the average persistence probability across all species for each of the 20 empirical mutualistic networks. Correlated (blue) and uncorrelated (red) habitat loss scenarios are compared. The three panels represent different landscape structures, highlighting how spatial configuration influences species persistence under varying habitat loss patterns.

More specifically, we find that nestedness of mutualistic networks has a positive effect on persistence probability across all landscapes and habitat loss patterns (Fig. 3, S3). However, the strength of this effect varies with both the landscape configuration and the pattern of habitat loss. In grid and random landscapes, the positive influence of nestedness weakens as habitat loss increases, with a more rapid decline under uncorrelated than correlated loss. Scale-free landscapes, in contrast, reveal a unique pattern: the effect of nestedness is relatively weak at both low and high levels of habitat loss but peaks at intermediate levels. Notably, the effect of nestedness remains stronger under uncorrelated loss in most fractions of habitat destruction. These results emphasize that the impact of the nested structure of communities on persistence is not static, but rather dynamically shaped by the landscape, the degree and the spatial pattern of habitat loss.

**Figure 3.**
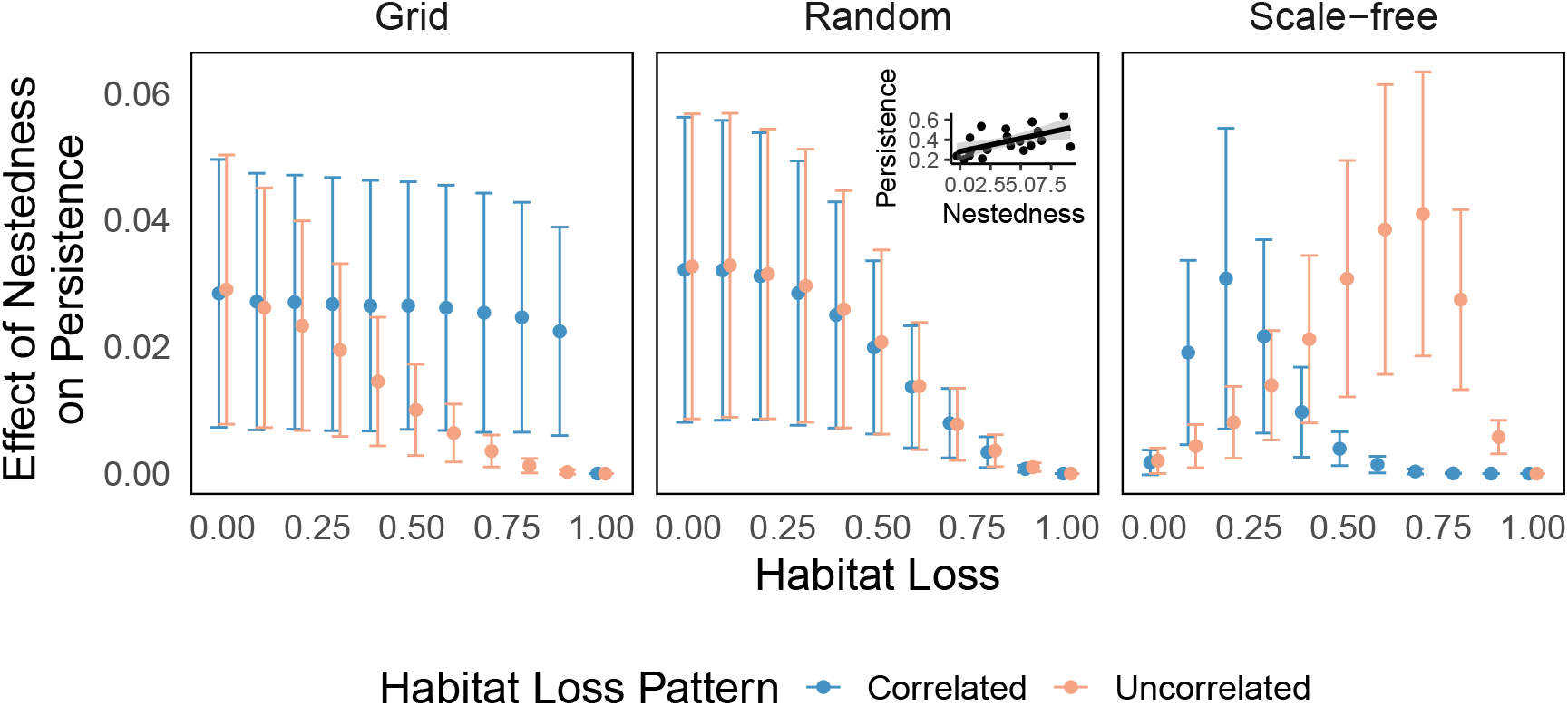
Effect of mutualistic community nestedness on average species persistence probability. Each point represents the estimated slope from a linear model capturing the relationship between nestedness and persistence probability at different habitat loss fractions. The lines indicate 95% confidence intervals, reflecting the uncertainty around the slope estimates. Correlated and uncorrelated habitat loss scenarios are shown in blue and red, respectively. The three panels correspond to different spatial network structures, highlighting how landscape configuration influences this relationship. The inset provides an example of the nestedness-persistence relationship for a specific habitat loss fraction (see Fig. S2 for additional scenarios across habitat loss fractions).

## Discussion

Our study investigates how habitat loss affects the persistence of species in mutualistic communities by examining the interplay between three key aspects: landscape configuration, community structure and habitat loss pattern. To quantify these effects, we performed simulations across a broad range of species extinction-to-colonization ratios and defined persistence probability as the fraction of simulations that yield nonzero regional species abundance. While previous studies have explored the three factors separately, our work highlights their combined effects, demonstrating that their interplay is critical for understanding species persistence.

Our findings show that the spatial configuration of habitat fragments shapes metacommunity persistence under habitat loss, with outcomes varying based on community structures and the pattern of habitat loss (Fig. 2). The trajectory of abundance loss, shaped by extinction-to-colonization dynamics, varies across different landscape configurations and is further modulated by the pattern of habitat loss—whether uncorrelated or correlated (Supplementary Fig. S1). Scale-free landscapes, characterized by a few highly connected hubs, show the highest persistence and extinction thresholds (Fig. S3) under spatially-uncorrelated habitat loss throughout the destruction process. These hubs serve as critical reservoirs that buffer species from extinction, maintaining metacommunity cohesion even when many peripheral patches are lost. However, this structural advantage becomes a liability under spatially-correlated habitat loss, particularly at low extinction to colonisation ratios (Fig. S4). The clustered removal of patches disproportionately impacts hub nodes, leading to cascading extinctions and rapid community collapse. In contrast, grid networks, with uniformly connected patches, are more resilient under correlated than uncorrelated habitat loss. Random networks show intermediate responses under both loss patterns. Thus, resilience is not solely dependent on landscape configuration, but also on how habitat is lost. These results broadly echo previous findings on the persistence of a single species (Gilarranz and Bascompte, 2012) and its responses to habitat loss (Heer et al., 2021; Liao et al., 2020).

We show that the structure of mutualistic communities also plays a pivotal role in species persistence under habitat loss. Specifically, we find that nestedness enhances persistence across all landscapes, thus offsetting some of the negative effects of habitat destruction (Fig. 3). This finding adds to the body of literature which demonstrates the importance of nestedness in promoting species coexistence, facilitating indirect interactions, and buffering communities against perturbations such as species loss or environmental fluctuations (Bascompte et al., 2003; Bastolla et al., 2009; Bhandary et al., 2023; Domínguez-Garcia et al., 2024; Lever et al., 2014; Thébault and Fontaine, 2010). However, we find that the positive effect of nestedness weakens with increasing habitat destruction. In homogeneous landscapes, this decline is gradual, whereas in scale-free networks, the effect of nestedness peaks at intermediate habitat loss fractions before sharply decreasing. This suggests that while nestedness can confer resilience in the early and intermediate stages of habitat loss, it may not be sufficient to prevent extinction cascades under more severe fragmentation. By explicitly incorporating both community structure and spatial landscape configuration, our study bridges a critical gap in understanding how mutualistic network architecture interacts with habitat loss to shape persistence outcomes.

From a conservation perspective, our findings suggest that protecting highly connected hub patches in heterogeneous landscapes or enhancing connectivity in more uniform landscapes can help maintain species persistence. Yet, the effectiveness of these strategies depends strongly on the type of habitat loss and community structure. Enhancing community nestedness, for example by protecting species which disproportionally contribute to the nested pattern of mutualistic interactions (Saavedra et al., 2011), may delay collapse. Yet, this strategy alone is insufficient under severe habitat degradation. Conservation planning should therefore integrate knowledge of both landscape and community structure, tailoring strategies to the specific nature of habitat loss. Such integrative approaches can enhance metacommunity stability, support early interventions before tipping points are reached, and provide a more reliable basis for preserving biodiversity in fragmented landscapes (Blüthgen and Staab, 2024; Gilarranz and Bascompte, 2012).

While our study offers valuable insights into how landscape configuration and mutualistic network structure influence species persistence under habitat loss, there are important aspects that merit further investigation. First, we assumed uniform habitat quality across all patches and did not account for species-specific habitat preferences. In real systems, variation in habitat quality—such as the presence of resource-rich or resource-poor patches—can strongly affect species persistence and the spatial stability of interactions (Ferraz et al., 2007; Hanski, 1999; Kremen et al., 2007). Incorporating spatial heterogeneity in habitat quality could refine our predictions, particularly for systems with specialized mutualisms. Second, our model treated all patches as equal in area, although empirical and theoretical work shows that patch size plays a critical role in shaping metapopulation dynamics and extinction thresholds (Fahrig, 2003; Hanski and Ovaskainen, 2003). Exploring scenarios where patch sizes vary—such as comparing a few large patches versus many small ones—could offer more nuanced insights into landscape design for conservation. Third, we focused solely on mutualistic interactions, yet real ecosystems include a mix of interaction types, including antagonistic and competitive, which may respond differently to habitat loss (Allesina and Tang, 2012; Bascompte and Jordano, 2007; Gawecka et al., 2022). Extending this framework to include mixed interaction types could help identify the role of mutualism within more complex, interaction-rich communities. Addressing these aspects in future work will enhance the generality of our framework and inform more effective conservation strategies in fragmented landscapes.

Empirical studies have demonstrated that habitat loss is a major driver of biodiversity decline, with cascading effects on ecosystem functions and species interactions. For example, long-term studies on pollination networks have shown that habitat fragmentation disrupts plant-pollinator mutualisms, reducing pollination success and leading to declines in plant reproductive output (Aguilar et al., 2006). Similarly, habitat loss in the Amazon has led to structural shifts in seed dispersal networks, altering connectivity patterns and species persistence (Galetti et al., 2013). These findings underscore the critical need to study how mutualistic networks respond to habitat loss within spatially structured landscapes. We show that the effects of habitat destruction on mutualistic communities vary across different landscape configurations and habitat loss patterns. However, the nested structure of mutualistic communities enhances species persistence, buffering some of the negative effects of habitat loss—and this buffering capacity itself changes depending on the landscape configuration. By integrating network theory with spatial metacommunity dynamics, our findings offer a framework for studying biodiversity loss and ecosystem collapse under ongoing habitat destruction. Understanding how mutualistic networks respond to habitat loss within different spatial contexts is crucial for developing targeted conservation strategies, particularly in landscapes undergoing rapid environmental change. These insights highlight the need for conservation planning that simultaneously considers the spatial landscape configuration and species interactions.

## Data Availability

Codes and data are available in a Github repository (https://github.com/subhendu-math/Habitat-loss-project.git).

## Acknowledgments

We thank June Monge Lorenzo and the members of Bascompte Lab for discussions. Funding was provided by SNSF (grant number 310030 197201 to JB), University of Zurich Postdoc Grant (grant number FK-22-114 to KAG), and Marie Skłodowska-Curie Actions Postdoctoral Fellowship (grant EP/Z000831/1 to KAG).

## Notes

https://github.com/subhendu-math/Habitat-loss-project

### Competing Interest Statement

The authors have declared no competing interest.

