## SUPPORTING INFORMATION for "Landscape configuration and community structure jointly determine the persistence of mutualists under habitat loss"

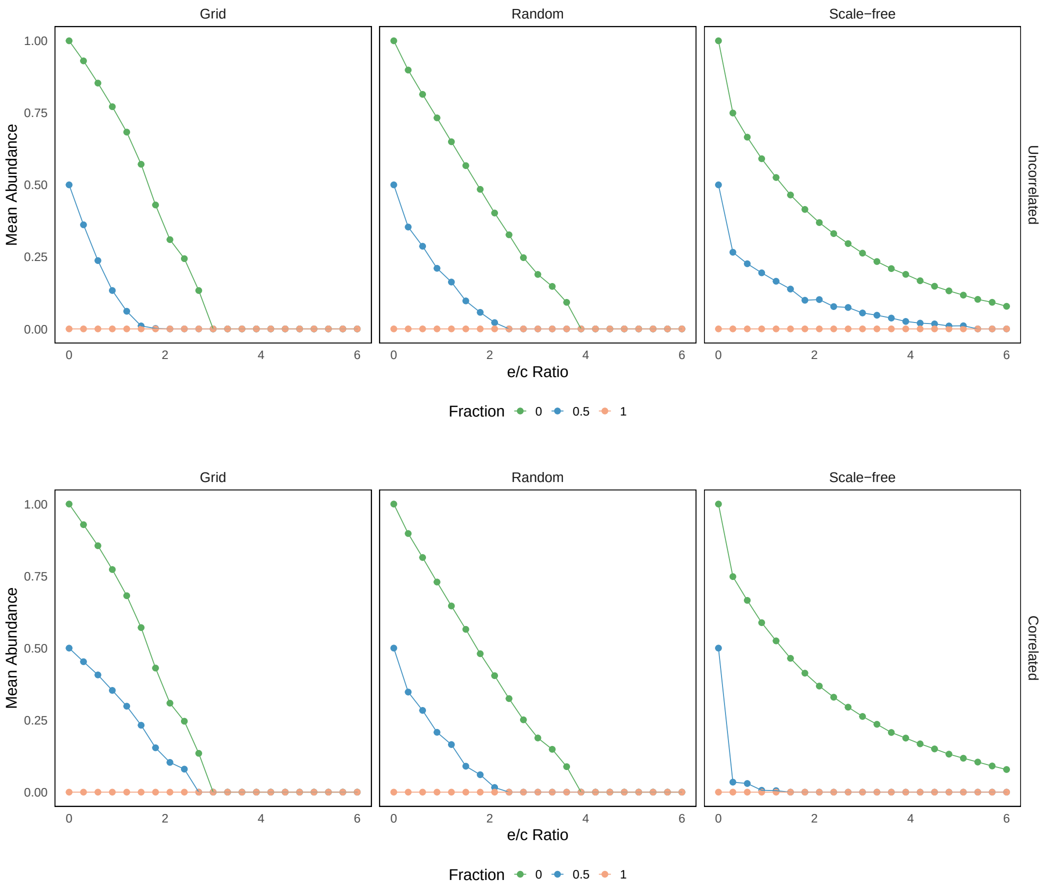


**Figure S1.** Mean species abundance as a function of increasing extinction-to-colonization ($e/c$) ratio across different landscape configurations. Each column represents a different landscape configuration (grid, random, scale-free), and each row corresponds to a habitat loss pattern: uncorrelated (top) and correlated (bottom). Within each subplot, colors represent different levels of habitat loss: low (green, 0), medium (blue, 0.5), and high (red, 1). For uncorrelated habitat loss with no degradation, scale-free networks maintain higher species abundance and do not reach extinction within the studied $e/c$ interval [0, 6], whereas in regular grids, mean abundance drops to extinction at lower $e/c$ ratios. In contrast, under correlated habitat loss, this pattern reverses beyond a threshold: at 50% habitat loss, mean abundance in scale-free landscapes collapses earlier than in regular grids. Results shown are for the mutualistic interaction network $M\_PL\_006$, assuming equal $e/c$ ratios for consumers and resources.

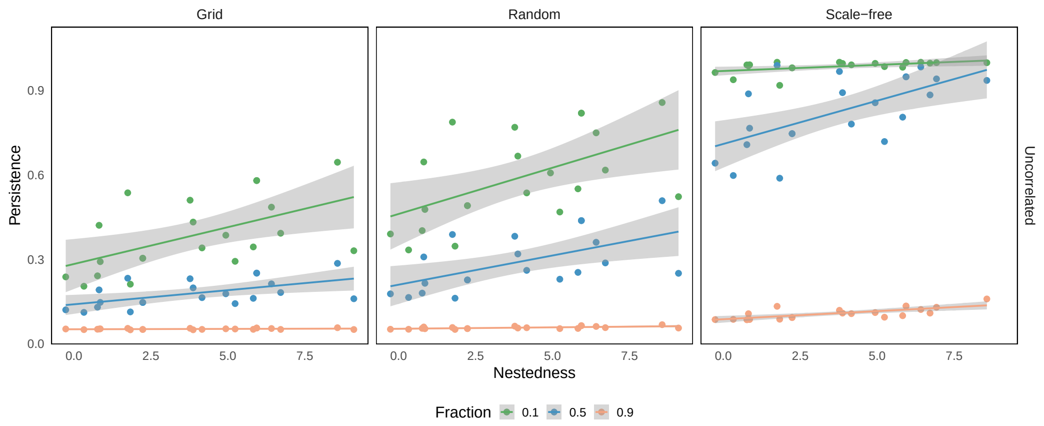


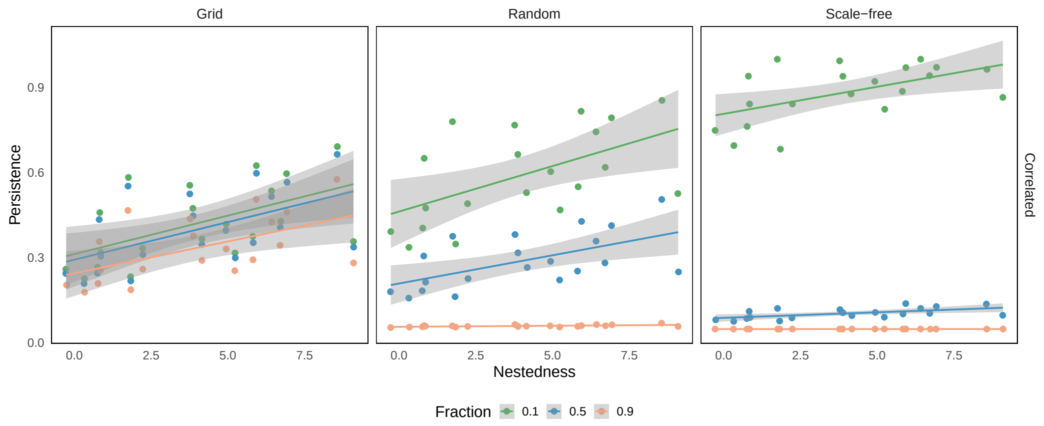


**Figure S2.** Effect of nestedness (expressed as a nest z-score) on species persistence probability under low (0.1), intermediate (0.5), and high (0.9) habitat loss fractions. Each subplot shows how this relationship varies across different landscape configurations (grid, random, scale-free), with color indicating the habitat loss fraction from low (green) to high (red). The top row presents results for uncorrelated habitat loss, while the bottom row shows correlated loss. Persistence increases with nestedness, but the slope of this relationship depends on habitat loss fraction, landscape configuration and habitat loss pattern.

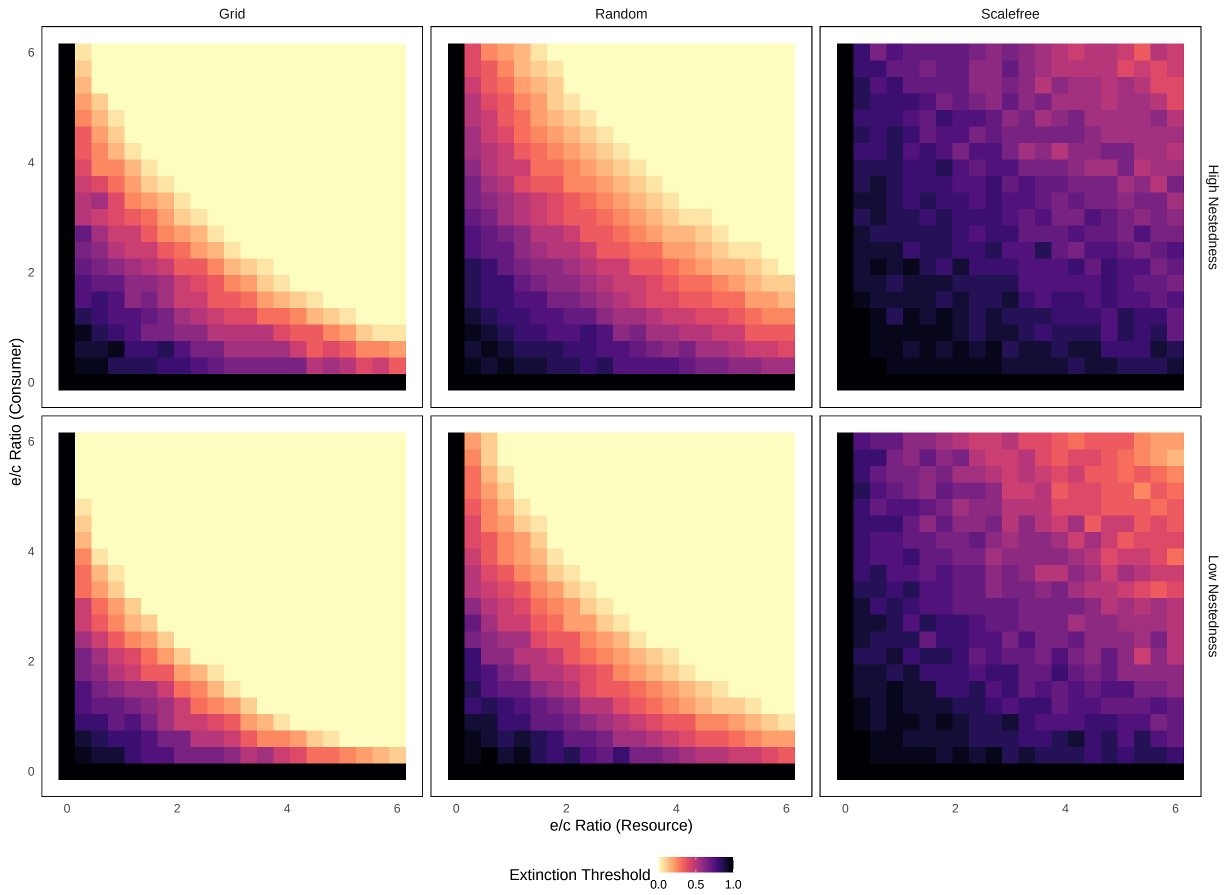


**Figure S3.** Effect of colonization–extinction ratio on extinction threshold under uncorrelated habitat loss. Colors show the extinction threshold habitat loss beyond which mean species abundance drops to zero, across varying extinction-to-colonization $(e/c)$ ratios. Each row corresponds to a different level of network nestedness: the top row for high-nestedness ($M\_PL\_006$) and the bottom row for low-nestedness ($M\_PL\_037$) mutualistic networks. Each column represents a different landscape configuration: grid, random, and scale-free. Grid landscapes show consistently lower extinction thresholds, indicating earlier collapse, whereas scale-free networks show higher thresholds (more persistence), with random networks in between. This pattern holds across both high and low nestedness levels. However, high-nested community structures demonstrate greater overall persistence than low-nested ones, highlighting the role of community structure in species extinction across all landscape configurations.

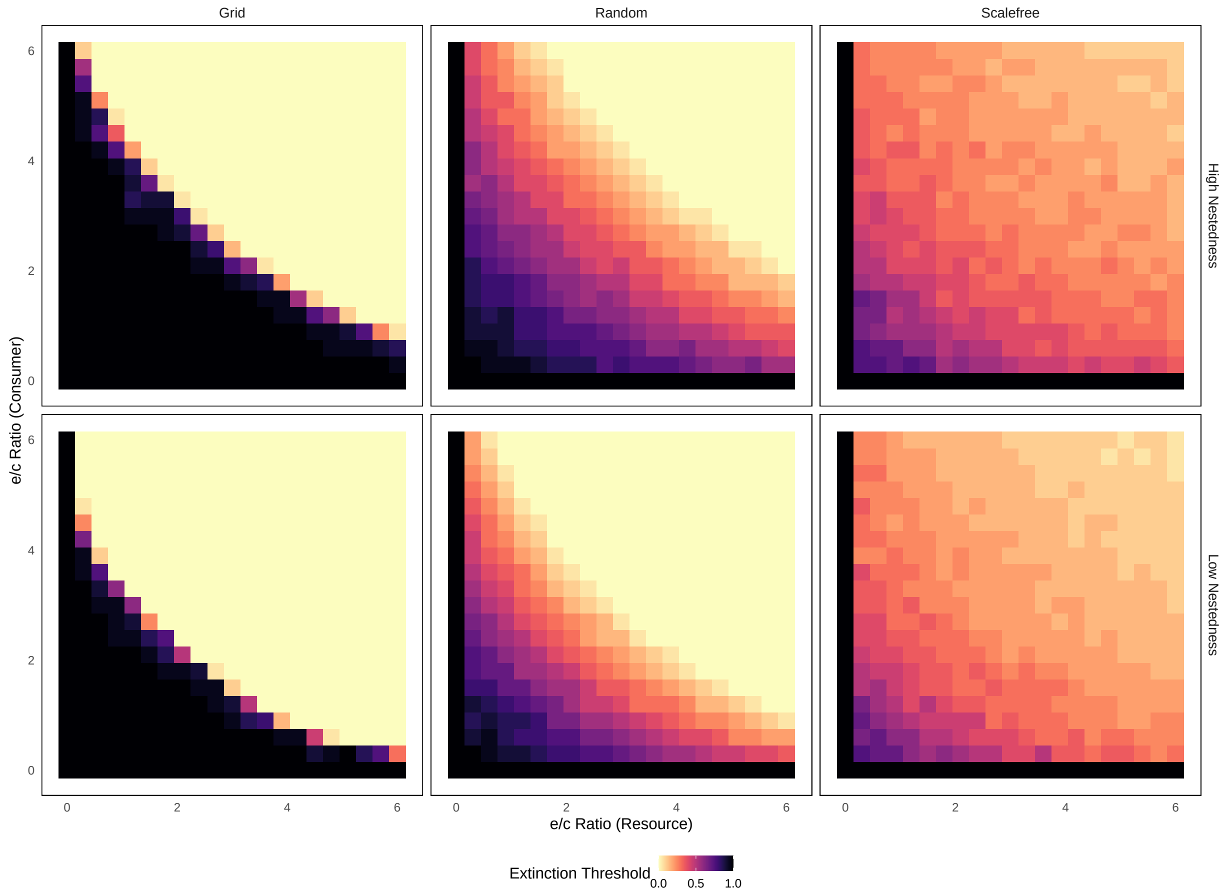


**Figure S4.** Effect of colonization–extinction ratio on extinction threshold under correlated habitat loss. Colors show the extinction threshold habitat loss beyond which mean species abundance drops to zero—across varying extinction-to-colonization ($e/c$) ratios. Each row corresponds to a different level of network nestedness: the top row for high-nestedness ($M\_PL\_006$) and the bottom row for low-nestedness ($M\_PL\_037$) mutualistic networks. Each column represents a different landscape configuration: grid, random, and scale-free. At low $e/c$ ratios, grid landscapes show higher extinction threshold, indicating delayed collapse, while scale-free networks show lower thresholds (earlier collapse), with random networks in between. However, at high $e/c$ ratios, the grid configuration exhibits earlier collapse, while scale-free networks show intermediate persistence. The transition in critical thresholds with increasing $e/c$ ratio is abrupt in grid landscapes (from black to yellow), but more gradual in random and scale-free networks. This trend holds for both high and low nestedness structures. However, high-nested community structures demonstrate greater overall persistence than low-nested ones, highlighting the role of community structure in species extinction across all landscape configurations.

**Table T1. Networks and their properties: Nestedness expressed as a z-score.** $S_A$ and $S_P$ stand for the number of Consumer(Animal) and Resource(Plant), respectively. We have considered plant–pollinator and seed dispersal networks as mutualistic interactions.

| Sl. No. | Network | $S_P$ | $S_A$ | Nestedness (z-score) |
|---|---|---|---|---|
| 1. | $M_PL_$006 | 17 | 61 | 9.0918 |
| 2. | $M_PL_$010 | 31 | 76 | 6.9250 |
| 3. | $M_PL_$025 | 13 | 44 | 4.9328 |
| 4. | $M_PL_$033 | 13 | 34 | 0.8164 |
| 5. | $M_PL_$036 | 10 | 12 | 0.3295 |
| 6. | $M_PL_$037 | 10 | 40 | -0.2671 |
| 7. | $M_PL_$039 | 17 | 51 | 2.2346 |
| 8. | $M_PL_$046 | 16 | 44 | 5.9359 |
| 9. | $M_PL_$051 | 14 | 90 | 5.2356 |
| 10. | $M_PL_$059 | 13 | 13 | 6.7148 |
| 11. | $M_SD_$002 | 31 | 9 | 3.8721 |
| 12. | $M_SD_$005 | 25 | 13 | 1.8319 |
| 13. | $M_SD_$007 | 72 | 7 | 4.1603 |
| 14. | $M_SD_$008 | 16 | 10 | 1.7470 |
| 15. | $M_SD_$010 | 50 | 14 | 3.7724 |
| 16. | $M_SD_$012 | 35 | 29 | 5.8259 |
| 17. | $M_SD_$014 | 16 | 17 | 6.4159 |
| 18. | $M_SD_$016 | 24 | 61 | 8.5606 |
| 19. | $M_SD_$025 | 7 | 6 | 0.7652 |
| 20. | $M_SD_$027 | 12 | 4 | 0.8540 |
